# Human Beta Cell Mass Expansion In Vivo With A Harmine and Exendin-4 Combination: Quantification and Visualization By iDISCO+ 3D Imaging

**DOI:** 10.1101/2020.07.24.220244

**Authors:** Carolina Rosselot, Alexandra Alvarsson, Peng Wang, Yansui Li, Kara Beliard, Geming Lu, Rosemary Li, Hongtao Liu, Virginia Gillespie, Nikolaos Tzavaras, Kunal Kumar, Robert J. DeVita, Andrew F. Stewart, Sarah A. Stanley, Adolfo Garcia-Ocaña

## Abstract

463 million people globally suffer from diabetes. The majority are deficient in insulin-producing pancreatic beta cells, although beta cells remain in most people with diabetes. Unfortunately, although many diabetes drugs exist, none is able to increase adult human beta cell numbers. Recently, small molecules that inhibit the kinase, DYRK1A, have been suggested to induce human beta cell replication *in vitro* and *in vivo* as assessed using proliferation markers, and this is enhanced by drugs that stimulate the GLP1 receptor (GLP1R) on beta cells. DYRK1A inhibitors also enhance human beta cell differentiation and function. However, it is unknown whether any drug can actually increase human beta cell mass *in vivo*, reflecting: 1) the intrinsic resistance of human beta cells to regeneration; and, 2) the current technical inability to accurately assess human beta cell mass *in vivo*. Here, we demonstrate for the first time that combining a DYRK1A inhibitor with a GLP1R agonist increases actual human beta cell numbers and overall mass *in vivo* by 400-700% in diabetic and non-diabetic mice over three months. We further describe a novel application of tissue-clearing and 3D imaging for quantification of human beta cell mass. These findings should be transformative for diabetes treatment.

## Introduction

Type 1 diabetes (T1D) results from autoimmune destruction of pancreatic beta cells, but most, and perhaps all people with T1D have residual beta cells at autopsy, and are able to secrete small amounts of insulin (1–5). Type 2 diabetes (T2D) is associated with insulin resistance together with a failure of beta cells to compensate by producing additional insulin. Beta cell mass is reduced in T2D by approximately 40-60%, attributable to genetic predisposition, metabolic stress, death and/or de-differentiation (6–11).

In spite of the clear need for beta cell regeneration, none of the myriad of drugs currently prescribed for millions of people with diabetes address the fundamental problem of beta cell deficiency. Some diabetes drugs, exemplified by the sulfonylurea, meglitinide, and GLP1RA classes, drive residual beta cells to produce additional insulin. Others, exemplified by metformin and the PPARγ classes, enhance insulin sensitivity. Others encourage glucose disposal by the kidney (the SGLT2 inhibitors) or impede glucose absorption by the intestine (the α-glucosidase inhibitors). When these drugs fail or are contraindicated, insulin itself is prescribed. Disappointingly, even in first-world countries, 70% of people with diabetes fail to achieve national and international guidelines for blood glucose control (12–14).

These considerations have provided impetus for development of alternate therapies, such as automated closed-loop insulin delivery systems with continuous glucose monitoring (“artificial pancreas”), whole pancreas transplant, pancreatic islet transplant, and transplant of human embryonic stem cells differentiated to beta cells (15–19). Each of these approaches represents a major advance in diabetes care, but none is scalable to the hundreds of millions of people with diabetes.

With the goal of developing an inexpensive, scalable treatment for millions of people with T1D and T2D that addresses the fundamental reduction in pancreatic beta cell mass and function, several groups have identified drugs that are able to induce residual human beta cells to proliferate with the hope of restoring normal functional beta cell mass (20–31). This is a formidable challenge, since adult human beta cells are terminally differentiated and recalcitrant to replication (32–33). In addition, human beta cells survive poorly in tissue culture, lasting only a matter of days before dying or de-differentiating (34). Nonetheless, this field has made rapid advances, most visibly in recent years with the identification and development of DYRK1A inhibitors, used alone or in combination with other classes of drugs such as the GLP1RA’s (20–31). In most cases, “replication” is evidenced by immunohistochemical markers of proliferation, exemplified by labeling indices for Ki67, BrdU or EdU, which increase from essentially zero to the 5-8% range (20–31). While this drug-induced human beta cell “proliferation” is exciting, it remains to be demonstrated that this translates into authentic, clinically relevant, increases in human beta cell numbers and mass.

DYRK1A inhibitors have an additional benefit: they are able to enhance human beta cell differentiation. For example, harmine and 5-IT increase expression of RNAs encoding beta cell specification factors and differentiation markers such as *PDX1, MAFA, MAFB, NKX6.1, PCSK1, GLP1R, SLC2A2*, accompanied by increased abundance of their cognate proteins in human beta cells, and by enhanced glucose-stimulated insulin secretion (GSIS) from human islets derived both from healthy donors as well as from those with T2D (20–22,26). Moreover, DYRK1A inhibitors enhance glycemic control and insulin secretion over the short term (two weeks) *in vivo* in immunodeficient diabetic mice transplanted with a marginal or therapeutically inadequate number of human islets (20–22).

Thus, overall, DYRK1A inhibitors, alone or in combination with GLP1RA’s, have remarkable beneficial effects on human beta cell proliferation and function in *in vitro* and in short term *in vivo* studies. However, three critical unmet challenges remain: 1) development of accurate quantitative tools to rigorously assess human beta cell mass in an *in vivo* model system; 2) demonstration that the regenerative effects of DYRK1A inhibitors, alone or in combination with GLP1RA’s translates into an authentic increase in human beta cell mass *in vivo*; and, 3) demonstration that this treatment can reverse diabetes over the long term (months). We address each of these challenges in this report.

## Results

### iDISCO^+^ Imaging Accurately Quantifies Human Beta Cell Mass *In Vivo*

We transplanted human pancreatic islets (300 islet equivalents, IEQ) into both right and left kidney capsules of immunodeficient RAG-1-deficient mice, followed by subcutaneous implantation of continuously infusing osmotic minipumps that deliver vehicle, harmine, the GLP1RA, exendin-4, or the combination of harmine plus exendin-4 for one week, four weeks, or three months, followed by *in vivo* perfusion and tissue fixation. One kidney was harvested for histology and immunohistochemistry, and the other for iDISCO+ tissue-clearing, immunolabeling for insulin and glucagon, light sheet microscopy and 3D image analysis to assess islet mass (**Fig. 1**). Volumetric graft measurements were validated by engrafting 100, 300, 500 or 1000 human IEQ into mouse kidneys followed by immediate harvesting of the transplanted kidney and measurement of beta cell and alpha cell mass (**Fig. 2, Supplemental Video 1**). These experiments demonstrated that actual measured beta and alpha cell volumes increased in an islet dose-related manner, and closely mimicked the predicted beta, alpha, and total cell volumes, providing confidence in the accuracy, precision and reproducibility of the iDISCO+ human islet graft measurement technique.

**Figure 1.**
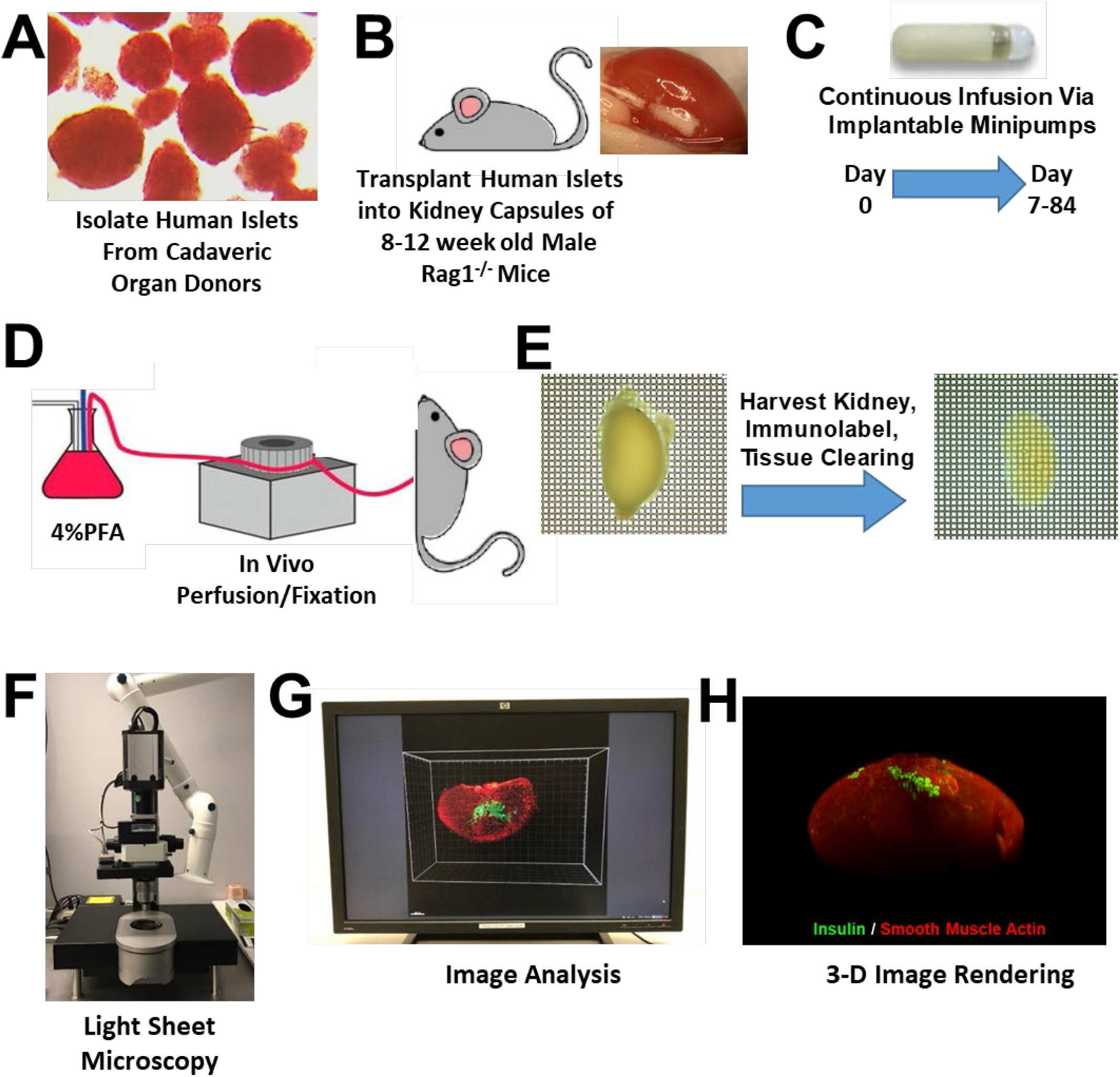
Eight Steps For Human Islet Transplant, Tissue-Clearing and 3D Imaging. Human islets (**A**) are transplanted under the renal capsule of both kidneys of immunodeficient mice (**B**) into which are implanted continuous infusion minipumps (**C**) containing vehicle (water), harmine, exendin-4 or the harmine-exendin-4 combination. After 1 week, 4 weeks, or 3 months, mice are perfusion-fixed (**D**), kidneys harvested and clarified by iDISCO+ and immunolabeled with insulin, glucagon or other antibodies (**E**), then examined by light-sheet microscopy (**F**), the Z-stacks from which are used to render 3D images using Imaris software (**G**), which can be used for 3D viewing and calculation of islet cell volume (**H**). See text and Materials and Methods for details.

**Figure 2.**
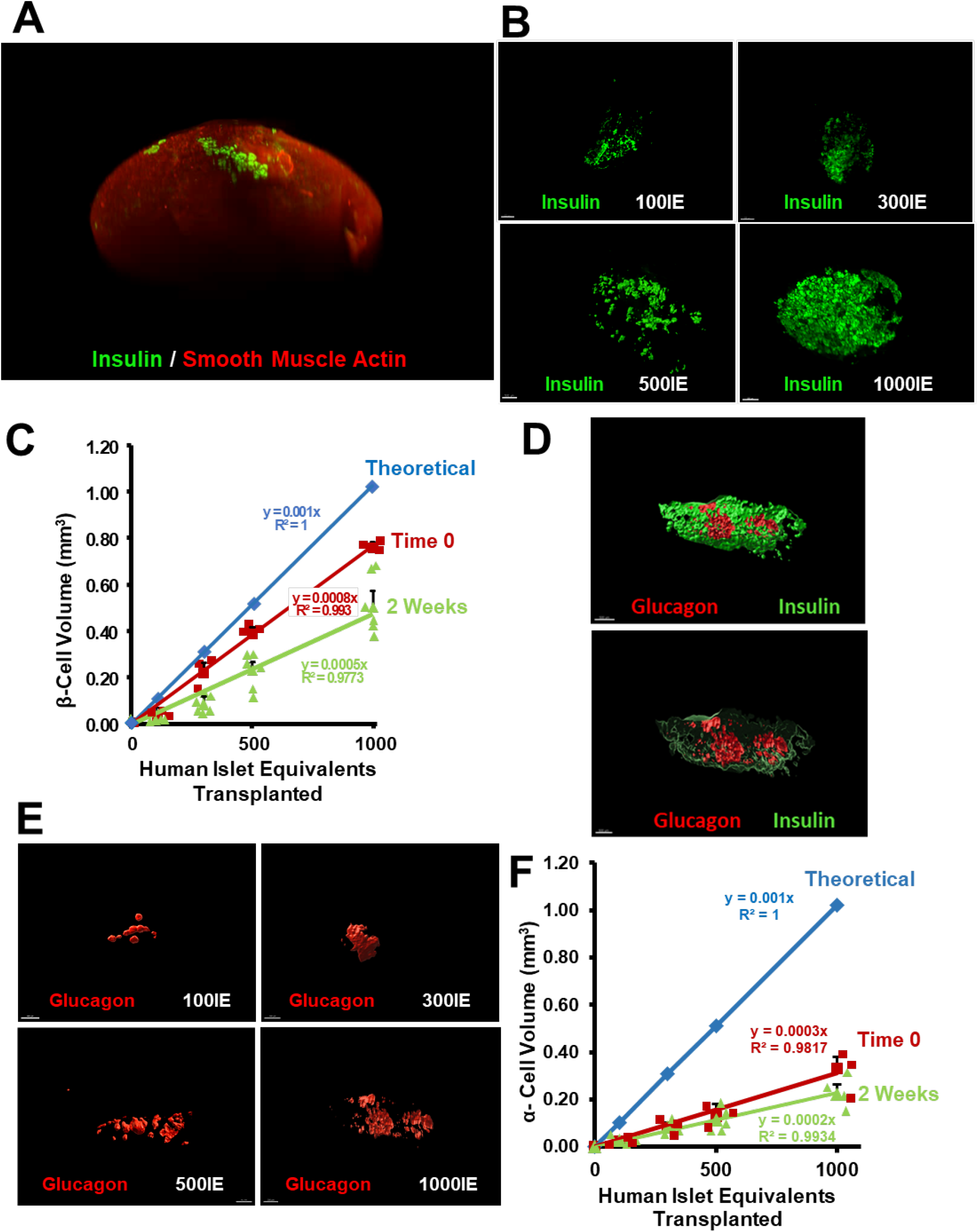
Quantification and Validation Of iDISCO+ Alpha and Beta Cell Imaging. (**A**) A 3-D image of beta cells in one human islet graft. Beta cells are immunolabeled in green, and smooth muscle actin immunolabeling in red reveals the entire mouse kidney. See 3-D Supplemental Video 1. (**B**) Examples of images of transplanted islets in progressively larger numbers of islet equivalents (IEQ) in immunodeficient mouse kidneys, and harvested immediately (insulin, green). (**C**) Mathematical validation of iDISCO+ quantification of human beta cell mass. The blue line represents the calculated, theoretical volume of 100, 300, 500 and 1000 intact 125 μm diameter IEQs, including all islet cell types, of which approximately 30-70% are beta cells or alpha cells. The red line represents the actual iDISCO+-measured volume of beta cells in 100, 300, 500 and 1000 actual human islets that were transplanted as in Panel B, after which the kidney was removed immediately and processed as described in Fig. 1. The green line is the same as the red line, except that the islet grafts were left in place for two weeks prior to harvesting and processing. Note that the actual volume of islets harvested immediately (red line) loosely follows theoretical blue line, but is approximately 35% lower reflecting the fact that not all cells in whole islets (blue line) are beta cells (red line). Note also that by two weeks (green line), beta cell volume in all islet preparations had declined by approximately 30%, likely reflecting additional beta cell death in the two-week interval between islet transplant and harvesting. Collectively, these data illustrate the accuracy, precision and reproducibility of iDISCO+ volume measurements. (**D-F**) This figure follows the design of panels B-C, but describes validation of the alpha cell volume measurements. (**D**) Human islet grafts were co-labeled for insulin (green) and glucagon (red) and subjected to processing as described in Fig 1. The upper image shows a view of a human islet graft as seen from the exterior. The lower image shows a cut through the center of the graft revealing alpha cells in the interior of the islet graft. (**E**) Static images of examples of alpha cells in grafts immunolabeled for glucagon in 100, 300, 500 and 1000 human IEQ. (**F**) See panel C for a detailed explanation. This panel is similar but assesses human alpha cell volumes in human islet grafts. Note that alpha cells comprise a smaller volume of total islet volume as expected, and that the sum of beta cell volumes in panel C plus the alpha cell volumes here are similar to total theoretical islet cell volume (blue lines) in both panels, reflecting the fact that the large majority of islet cells are either alpha or beta cells.

### Harmine and Exendin-4 Dose Selection

To determine an optimal dose of harmine for infusion, we first implanted minipumps into mice delivering vehicle or progressively increasing doses of harmine for one week, followed by analysis of mouse beta cell and alpha cell proliferation, as defined by Ki67 immunolabeling. As shown in **Supplemental Fig. 1**, harmine infused at 10 mg/kg/day show a marked significant increase in alpha and beta cell proliferation, similar to a control group receiving harmine by daily i.p. injection at 10 mg/kg (20–22). Importantly, we observed that infusion of harmine at 3 mg/kg/day led to a significant increase in human beta cell proliferation, but no increase in alpha cell proliferation. Thus, this dose of harmine, alone or in combination with the infusion of the previously reported dose of exendin-4 (0.1 mg/kg/day) (33), was selected in subsequent studies.

### A 700% Increase In Human Beta Cell Mass In Nondiabetic Mice

We next assessed the effects of vehicle, harmine, exendin-4, or the harmine-exendin-4 combination on beta cell volume, proliferation and cell death. At one-week post-transplant, human beta cell volume was 50% higher in the harmine-exendin-4 group compared with the vehicle group (**Fig. 3A,B**), associated with a striking increase in beta cell proliferation (Ki67), and a corresponding reduction in beta cell labeling for TUNEL, a marker of cell death (**Fig. 3C,D**). Taken together, these observations suggest that the increase in beta cell volume from the drug combination reflects components both of enhanced beta cell proliferation and survival.

**Figure 3.**
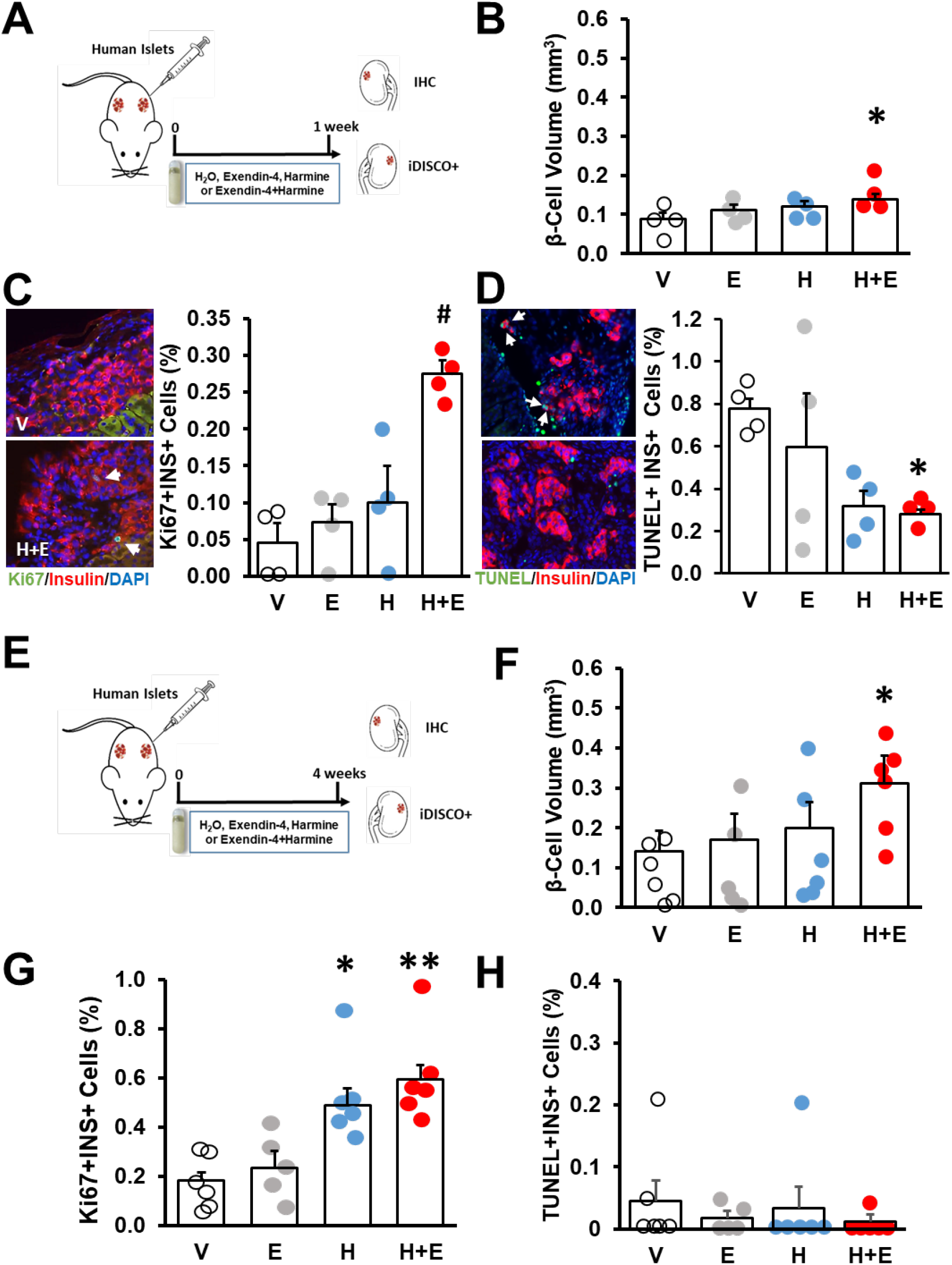
Effects Of Vehicle, Harmine, Exendin-4 and The Combination On Human Beta and Alpha Cells Following One Week and Four Weeks Of Treatment. (**A**) The one-week study protocol. Islets are transplanted, minipumps implanted and one week later both kidneys are harvested for immunohistochemical and iDISCO+ analysis. (**B**) Human beta cell volume in the four groups of mice at one week. Dots represent individual islet grafts from four different human islet donors in four different mice. (**C**) Examples of Ki67 immunolabeling in vehicle and harmine plus exendin-4 groups. Arrows indicate Ki67+ insulin+ cells. Labeling index of Ki67 in human beta cells in grafts from the four different donors. (**D**) Examples of labeling for TUNEL, a marker for cell death, in human beta cells from grafts in mice treated with vehicle, or harmine plus exendin-4. Arrows indicate TUNEL+ insulin+ cells. Quantification of TUNEL labeling in beta cells of islet grafts from each of the islet preparations. (**E**) The four week study protocol. Islets are transplanted, minipumps implanted and four weeks later both kidneys are harvested for immunohistochemical and iDISCO+ analysis. (**F**) Human beta cell volume in the four groups of mice at four weeks. Dots represent individual islet grafts from five to six different human islet donors in five to six different mice. (**G**) Labeling index of Ki67 in human beta cells in grafts from five to six different donors. (**H**) Quantification of TUNEL labeling in beta cells of islet grafts from each of the islet preparations. In all panels, error bars indicate mean ± SEM, * indicates p<0.05 vs V; ** indicates p<0.05 vs V and E groups; and # indicates p<0.05 vs V, E and H groups.

After four weeks of treatment, beta cell mass was comparable in the vehicle, exendin-4 alone and harmine alone groups (0.1-0.2 mm^3^), but significantly greater (0.31 mm^3^) in the combination group (**Fig. 3E,F),** presumably reflecting the increased survival observed in the first days after transplantation (**Fig. 3D**) together with a continued increase in human beta cell replication in the harmine-exendin-4 treated mice (**Fig. 3G,H**).

Based on these encouraging results, we next extended these studies to three months in a third cohort of mice. Following three months of treatment, average beta cell volume was 0.1 mm^3^ in the vehicle and exendin-4 groups, 0.34 mm^3^ in the harmine group, and 0.74 mm^3^ in the harmine-exendin-4 group (**Fig. 4A-C, Supplemental Videos 2 and 3**). Compared to the vehicle and exendin-4 groups, human beta cell proliferation was 2-3-fold higher in the harmine alone and the harmine-exendin-4 groups (**Fig. 4D**) while beta cell death was essentially absent in all groups (**Fig. 4E**). Beta cell size at 3 months was comparable among all four groups (**Fig. 4F**).

**Figure 4.**
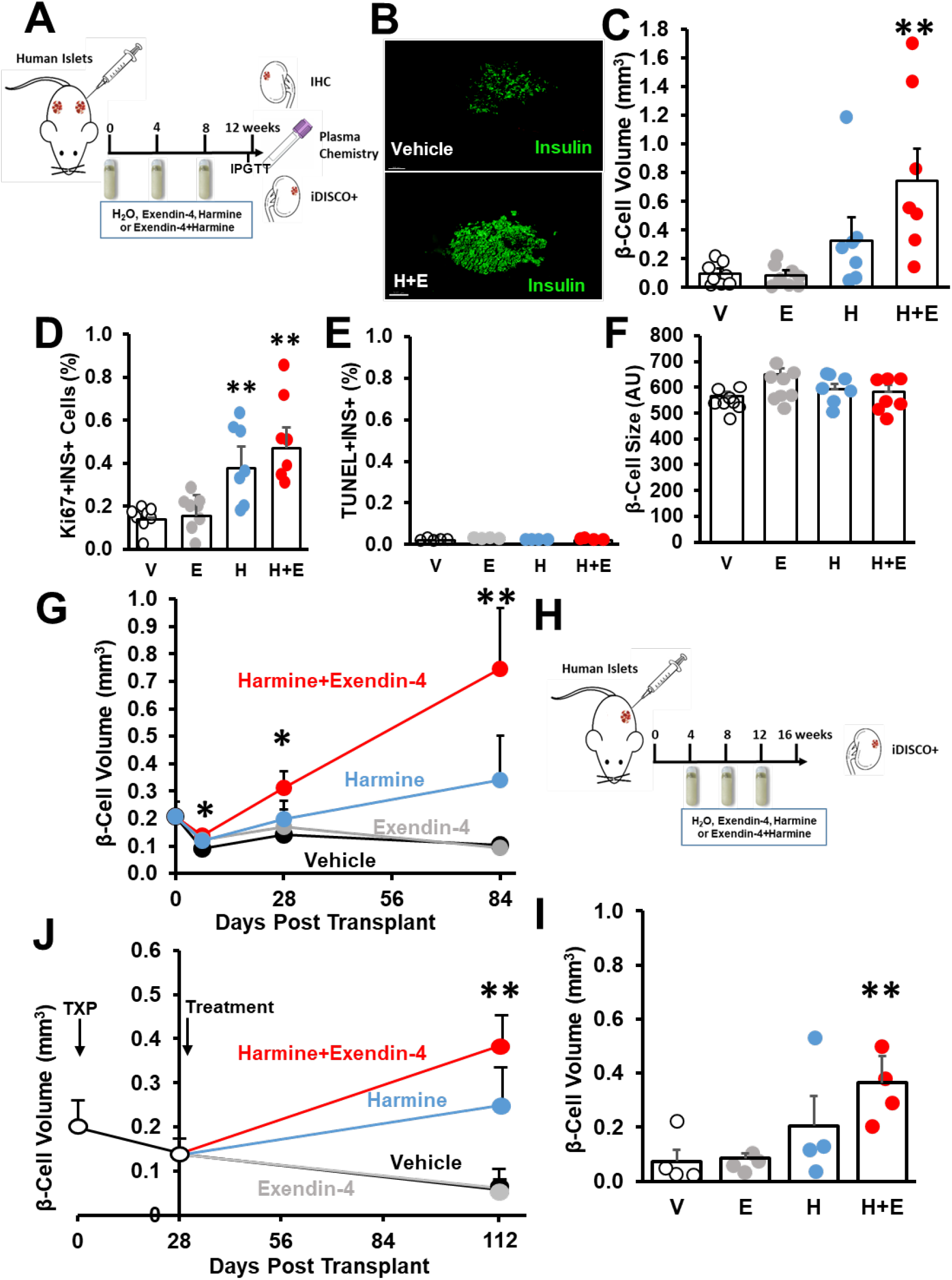
Effects Of Three Months Of Harmine and/or Exendin-4 On Human Beta Cell Mass and Function. (**A**) The three-month study protocol. Four-week minipumps are implanted and replaced at one month and two months to provide three months of continuous infusion. At month 3, both kidneys are harvested and blood is obtained. (**B**) Static images of kidney grafts treated for three months with vehicle (top) or the harmine plus exendin-4 (bottom) for three months. See 3D Supplemental Videos 2 and 3. (**C**) Human beta cell volume in the four groups of mice at three months. Dots represent individual islet grafts from seven to eight different human islet donors in seven to eight different mice. (**D**) Labeling index of Ki67 in human beta cells in grafts from each of the islet preparations. (**E**) Quantification of TUNEL labeling in beta cells of islet grafts from each of the islet preparations. (**F**) Beta cell size assessed in islet grafts stained for insulin as previously described (56), and the size of a minimum of 300 beta cells was quantified per section using ImageJ in each of the four groups. (**G**) Summary of the changes in human beta cell volume at all time points in the four treatment groups. Note that after an initial decline in each group, harmine alone, and the harmine plus exendin-4 combination progressively increases beta cell mass by 3-fold and 7-fold, respectively, as compared to vehicle or exendin-4. (**H**) The four week delay after transplant before three-month treatment study protocol. Islet are transplanted and 4 weeks later minipumps are implanted and replaced at one month and two months to provide three months of continuous infusion. At month 3 of treatment, kidneys are harvested for iDISCO+ analysis. (**I**) Human beta cell volume in the four groups of mice at three months. (**J**) Summary of the changes in human beta cell volume at three months in the four treatment groups. Values at time 0 and 28 days are from panel G vehicle-treated mice and serve as a starting point or reference for the changes induced by three months of treatment. In all panels, error bars indicate mean ± SEM, * indicates p<0.05 vs V, ** indicates p<0.05 vs V and E groups.

The collective results over time are summarized in **Fig. 4G**, which illustrates several points. First, beta cell mass declined in the first week post-transplant in all groups, as expected (34,35), but this was less in the harmine-exendin-4 group, presumably reflecting increased beta cell survival and proliferation at this early time point (**Fig. 3B-D**). Beta cell mass then progressively increased over the ensuing three months in the harmine and harmine-exendin-4 groups, at which point beta cell mass was 3-fold greater than at the time of initial transplant, and 7-fold greater, respectively, as compared to the vehicle and exendin-4 groups. This represents the first demonstration that it is possible to markedly increase human beta cell mass *in vivo* by any drug treatment.

Since beta cell transplantation in mouse and man is associated with substantial cell death in the first 24 hours post-transplant (34,35), and since harmine alone or in combination with exendin-4 attenuated cell death (**Fig. 3D**), we wondered to what extent the early beneficial survival effects of harmine alone or the harmine-exendin-4 combination contributed to the overall increase in beta cell mass at three months (**Fig. 4G**). To address this question, we repeated the three month studies with a modification (**Fig. 4H,J**): we transplanted human islets in four groups of animals as in prior studies, but delayed drug treatment for four weeks, to allow equal human islet engraftment and vascularization in all four groups, so that they would begin treatment with the same initial beta cell mass. We then analyzed beta cell volume in human islet grafts following three months of treatment. Following three months of treatment, average beta cell volume was 0.07-0.08 mm^3^ in the vehicle and exendin-4 groups, 0.20 mm^3^ in the harmine group, and 0.36 mm^3^ in the harmine-exendin-4 group (**Fig. 4I**). These results bring out two important interpretations: 1) the early (one week) human beta cell pro-survival effect and enhanced beta cell mass induced by the combination (**Fig. 3A-D)** play an important role in the ultimate increase in beta cell mass observed at three months of treatment, since the same treatment initiated one month post-transplant leads to a more moderate beta cell mass expansion (0.74 vs. 0.34 mm^3^); and, 2) the three-fold increase in beta cell mass in this context (**Fig. 4I**) correlated with the three-fold increase in human beta cell proliferation observed with the combination compared with vehicle-treated mice (**Fig. 4D**). These observations suggest that both beta cell proliferation as well as enhanced survival contribute to the overall 7-fold increase in beta cell mass observed in **Fig 4C,G**.

### Enhanced Glucose Tolerance With Harmine-Exendin-4 Treatment

Following three months of treatment, body weights were comparable among the four groups, but blood glucose was lower in all three drug treatment groups, and plasma human insulin, but not plasma glucagon, was elevated in the harmine-exendin-4 group (**Figs. 5A-D**). Glycemic responses to intraperitoneal glucose administration were improved in all three treatment groups as compared to the vehicle group, but were most improved in the harmine-exendin-4 group (**Fig. 5E**). Immunohistochemical evaluation of key beta cell transcription factors (PDX1, NKX6.1 and MAFA) in the human islet grafts indicated that these were maintained and comparable in all four groups (**Fig. 5F**). Collectively, these findings represent the first demonstration that it is possible to markedly increase human beta cell mass while also retaining or improving function over the long term *in vivo*.

**Figure 5.**
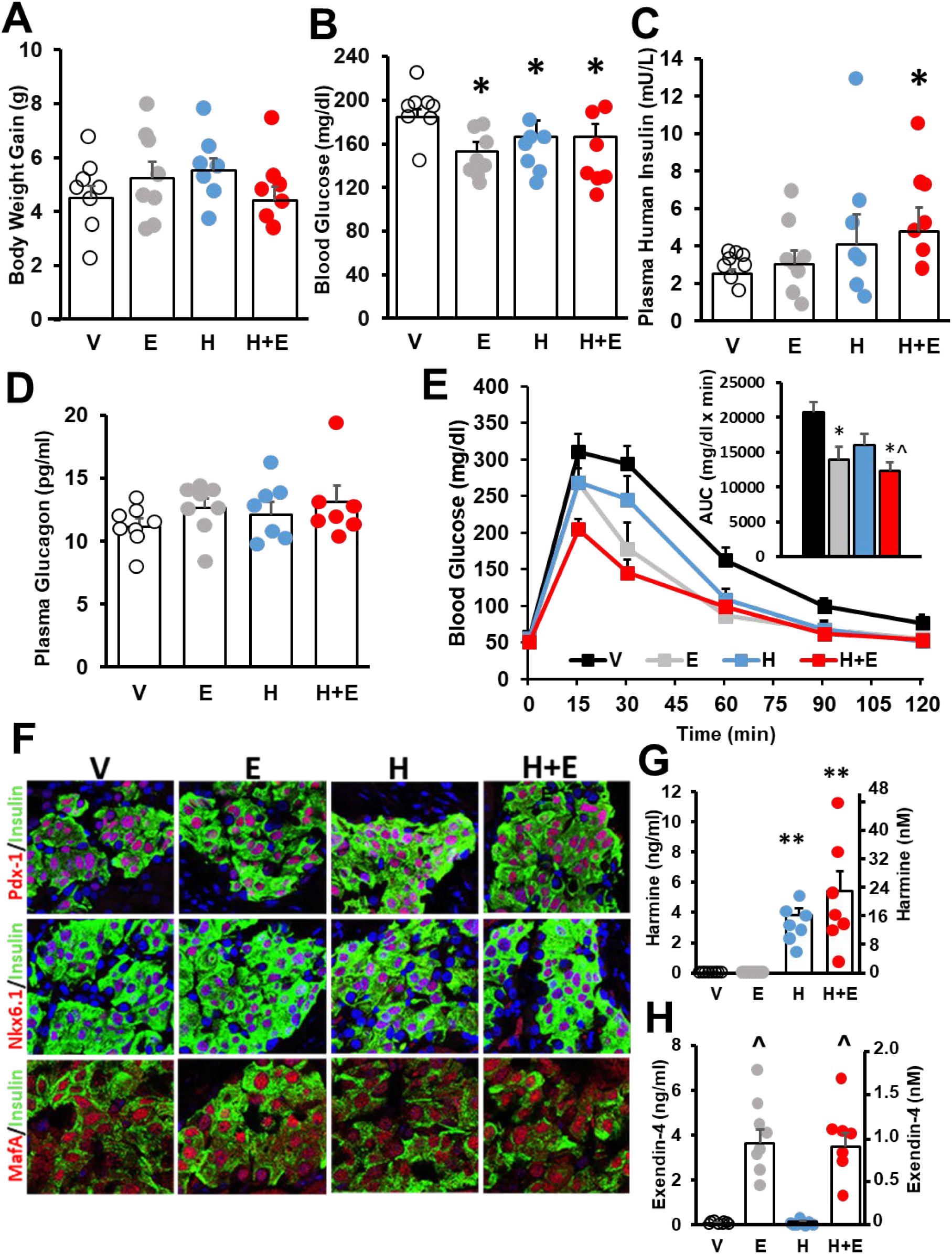
Effects On Human Beta Cell Function At Three Months. (**A**) Body weight gain, (**B**) random blood glucose, (**C**) plasma human insulin and (**D**) plasma glucagon in the four groups at the end of three-month treatment. (**E**) Intraperitoneal glucose testing in the four groups. Inset shows area under the curve for each group. (**F**) Human beta cells in islet grafts remained differentiated in the four groups as reflected in equivalent expression of PDX1, NKX6.1 and MAFA transcription factors. (**G**) Plasma harmine concentrations and (**H**) plasma exendin-4 concentrations in the four groups at month 3. In all panels, error bars indicate mean ± SEM, * indicates p<0.05 vs V, ** indicates p<0.05 vs V and E groups, ^ indicates p<0.05 vs V and H groups.

Finally, we assessed steady-state circulating harmine and exendin-4 concentrations in mice continuously infused with harmine, exendin-4 or the combination for three months (**Fig. 5G,H**). Mean harmine and exendin-4 concentrations were approximately 4-5 ng/ml (~16-20 nM) and 4 ng/ml (~1 nM), respectively.

### Harmine and Exendin-4 Effects On The Alpha Cell

Harmine and other DYRK1A inhibitors can activate rodent and human alpha cell proliferation *in vitro* (7–9,13) and *in vivo* (**Supplemental Fig. 1**). Guided by these preliminary data, we selected harmine doses below those associated with rodent alpha cell proliferation for the studies in **Figs 3,4**. We next assessed human alpha cell proliferation and volume *in vivo* in the same mice transplanted with human islets in **Figs 3,4**. Harmine and the harmine-exendin-4 combination caused a small, but non-significant increase in alpha cell Ki67 immunolabeling at one and four weeks that became significant at three months (**Figs 6A-D**). However, the rates of alpha cell proliferation were low (0.2-0.3%). Alpha cell death was negligible and not significantly different among the groups at three-months (**Fig. 6E**).

**Figure 6.**
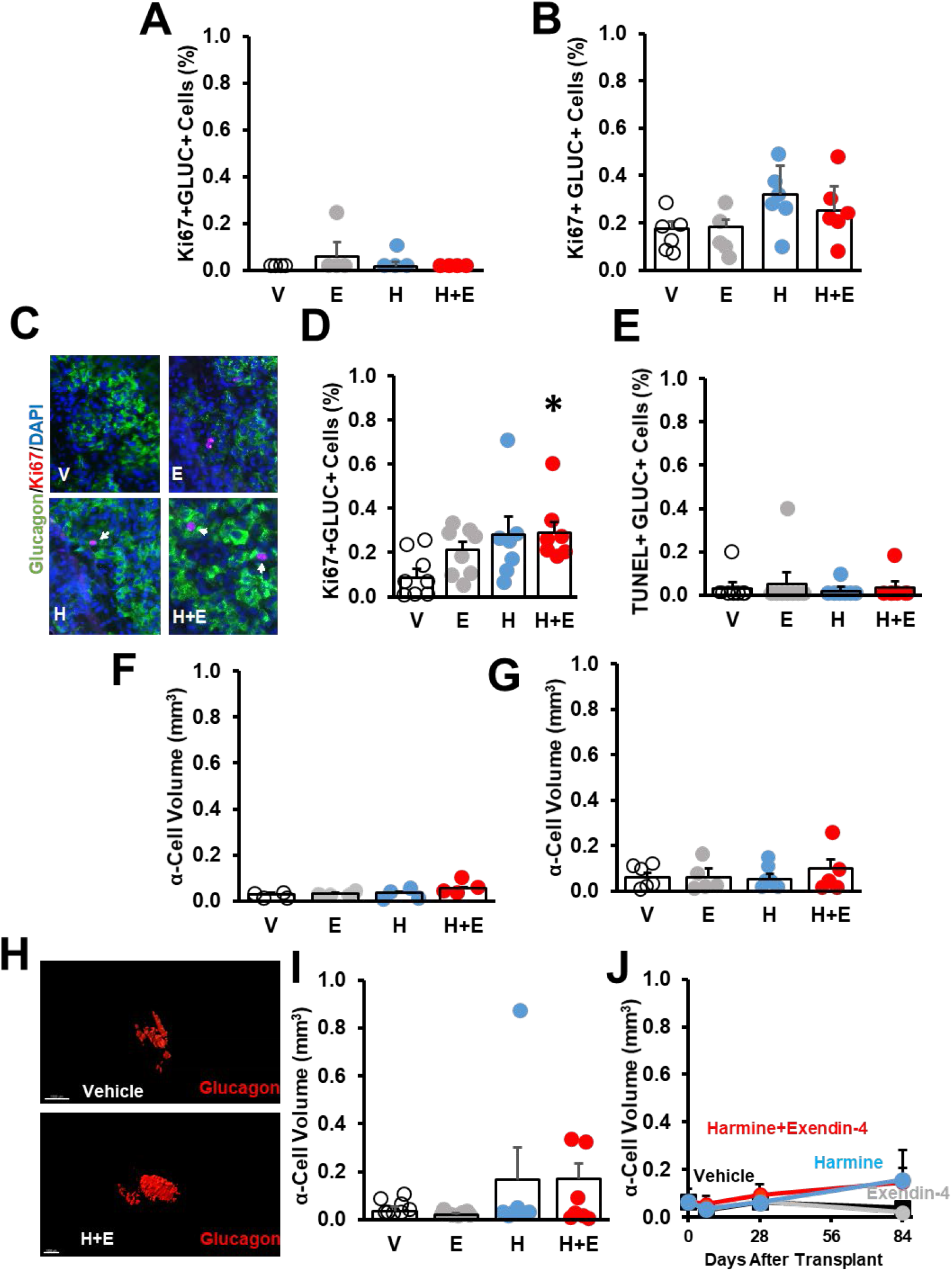
Effects On Human Alpha Cell Replication, Death and Mass At One Week, Four Weeks and Three Months. (**A**) Labeling index of Ki67 in human alpha cells in grafts from four different donors at one week. (**B**) Labeling index of Ki67 in human alpha cells in grafts from five to six different donors at four weeks. (**C**) Examples of Ki67 immunolabeling in the four groups. Arrows indicate Ki67+ insulin+ cells. (**D**) Labeling index of Ki67 in human beta cells in grafts from the seven to eight different donors at three months. (**E**) Quantification of TUNEL labeling in beta cells of islet grafts from each of the islet preparations at three months. Quantification of alpha cell volume in the same samples as in Fig. 3 at (**F**) one week and (**G**) four weeks. (**H**) Examples of alpha cell 3D images from vehicle and harmine plus exendin-4 treated mice at 3 months. (**I**) Human alpha cell volume in the same samples of the four groups of mice at three months represented in Fig. 4. (**J**) Summary of changes in alpha cell mass at each time point. Alpha cell mass appears to trend upwards in the harmine and harmine plus exenatide groups, but does not achieve significance. In all panels, error bars indicate mean ± SEM. * indicates p<0.05 vs V.

We next performed iDISCO+ tissue clearing and immunolabeling for glucagon on the same one-week, four-week and three-month kidney samples used for insulin/beta cell labeling (**Figs. 3,4**). As shown in **Figs. 6F-J**, despite the modest increase in alpha cell proliferation at three months, and despite the increase in beta cell mass in the same samples, there were no significant changes in alpha cell mass with any treatment at any time point. Thus, beta cell mass expansion can be dissociated from alpha cell mass changes.

### Three Month Safety Of The Harmine-Exendin-4 Combination

Harmine and exendin-4, if given at sufficiently high doses, may have undesired effects on cells and tissues outside the beta cell, since DYRK1A is ubiquitous, and GLP1R expression, although far more limited, is not exclusive to the beta cell (36,37). Accordingly, we explored proliferation as well as tissue histology in a wide range of tissues in the same RAG-1-deficient mice that had received the human islet transplant (**Fig. 4A-G**), including liver, spleen, intestine, adipose tissue, lung, heart, kidney, exocrine and endocrine pancreas, and brain (**Figs. 7–9**). In brief, while proliferation was observed in endogenous mouse beta cells as expected (**Fig. 7E**), no evidence of abnormal proliferation or tissue histology was observed in any other organ, nor were abnormal posturing, feeding or grooming behaviors observed. Equal numbers of mice in both groups did display innate immune cell abnormalities in the lung, presumably reflecting their immunocompromised state, but there were no differences between the groups (**Fig. 9**).

**Figure 7.**
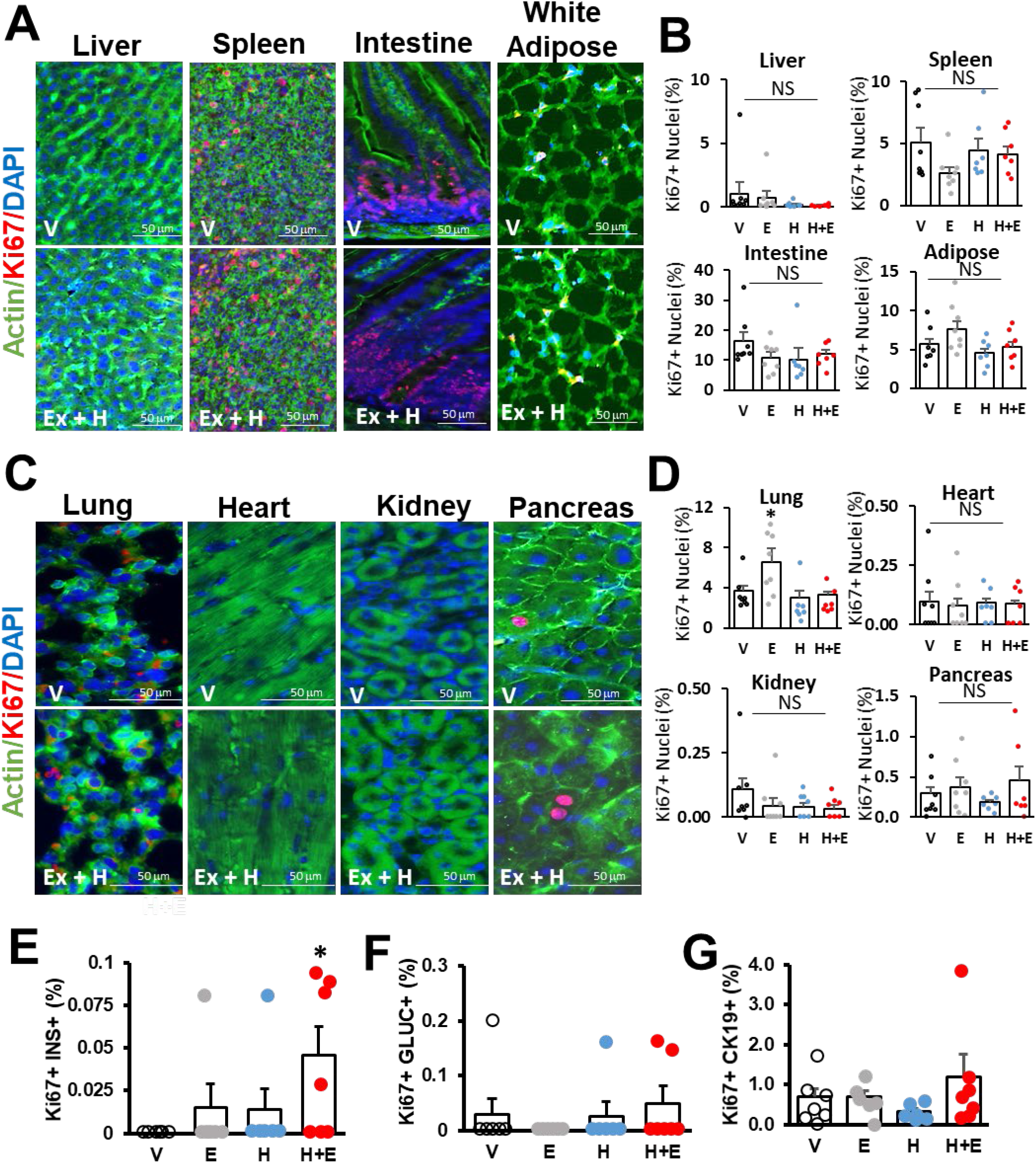
Assessment Of Off-Target Proliferation In RAG-1 Mouse Tissues. (**A-D**) Following three months of treatment, Ki67 immunolabeling was assessed and quantified in mouse liver, spleen, intestine, white adipose tissue, lung, heart, kidney and pancreas. (**E, F** and **G**) Endogenous mouse beta cell, alpha cell and ductal cell proliferation. As expected^7-9^, harmine plus exendin-4 induced beta cell proliferation in endogenous mouse islets (*p=0.05 vs. V) as in human beta cells (Fig 4D). No significant differences among groups were observed in the Ki67 labeling index in mouse alpha cells or ductal cells. In all panels, error bars indicate mean ± SEM.

**Figure 8.**
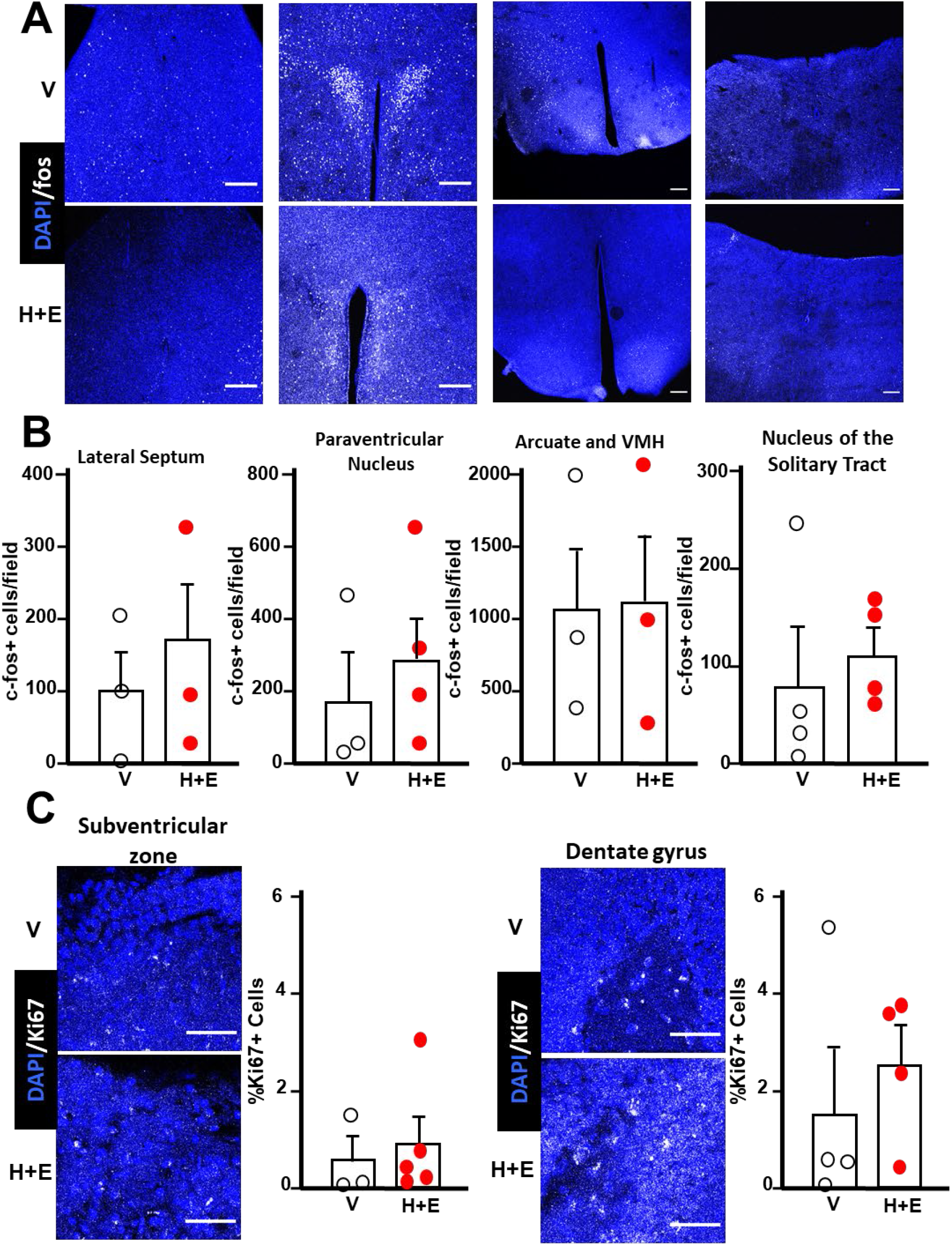
c-Fos and Ki67 Expression In The Brain. Brains from a subset of the same groups of mice in Fig. 4 and 7 were harvested. (**A**) and (**B**), cells in CNS nuclei known to have GLP1 receptors (lateral septum, paraventricular, arcuate, ventromedial hypothalamus, nucleus of the solitary tract) were examined for c-Fos activation. **C.** Cells in the subventricular zone and dentate gyrus, also known to possess GLP1 receptors and with reported cell proliferation were examined for proliferation using Ki67 as a proliferation marker. Overall, no significant increases in c-Fos or Ki-67 expression were observed at any site. In all panels, error bars indicate mean ± SEM.

**Figure 9.**
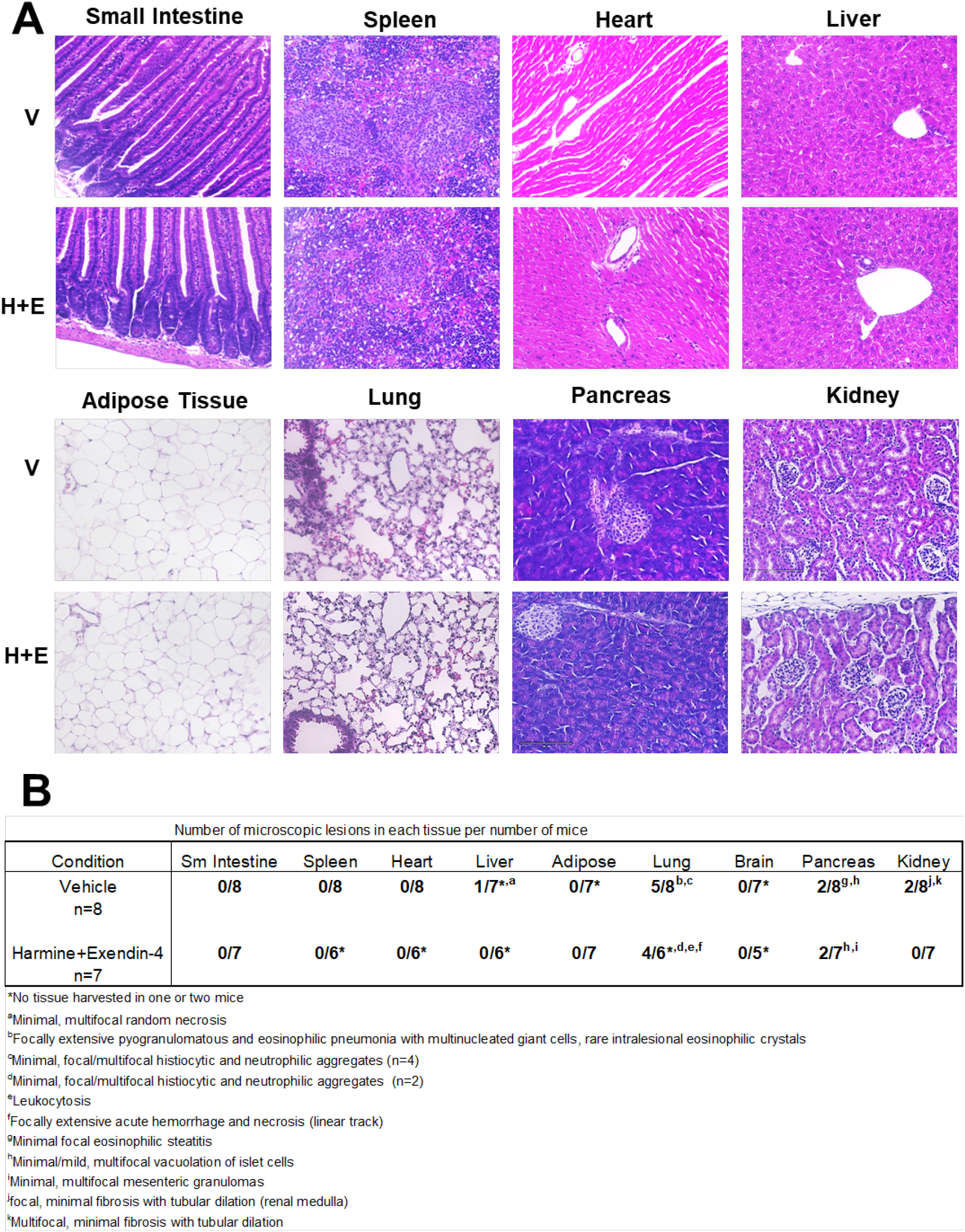
Histology In Tissues From RAG-1 Mice Treated With Vehicle or The Harmine-Exendin-4 Combination. At the conclusion of the three-month treatment protocol, the tissues described here were harvested, formalin-fixed, paraffin-embedded, sectioned, stained for hematoxylin and eosin, and sent for review by a veterinary pathologist in a double-blinded manner. (**A**) Examples of the 8 tissues examined from animals treated with vehicle (V) or harmine + exendin-4 (H+E). (**B**) A table showing the numbers of animals examined (7 or 8 for each organ/tissue), the number of animals with observed abnormalities, and footnotes describing the abnormalities observed. Several mice in both groups had inflammatory abnormalities in the lung, which are characteristic of RAG-1 mice. The important point is that there were no differences between the treated and control groups.

### Harmine-Exendin-4 Combination Treatment Increases Human Beta Cell Mass and Improves Glucose Homeostasis In Streptozotocin (STZ)-Induced Diabetic Mice

To assess the effect of continuous treatment with harmine-exendin-4 on human beta cell mass and glucose homeostasis in a conventional model of diabetes, we performed long term, three month experiments in STZ-induced diabetic immunodeficient RAG1-deficient mice transplanted with marginal or therapeutically inadequate number of human islets (**Fig. 10A**). Diabetic mice transplanted with human islets and treated with harmine-exendin-4 combination displayed a rapid, significant and sustained normalization of blood glucose concentrations that was maintained for the entire three months of treatment, and superior to the vehicle and single drug treatment groups (**Fig. 10B,C**). This correlated with a statistically significant increase in plasma human insulin in the harmine-exendin-4 combination treatment group (**Fig. 10D**). Glucose tolerance at three months was also improved most significantly in the harmine-exendin-4 treated mice compared with the vehicle and harmine-alone groups (**Fig. 10E**). Collectively, these data indicate that harmine-exendin-4 treatment provides significant, and sustained improvement in glucose homeostasis not only in euglycemic models (**Fig. 5B-E**), but also in a diabetic model of T1D.

**Figure 10.**
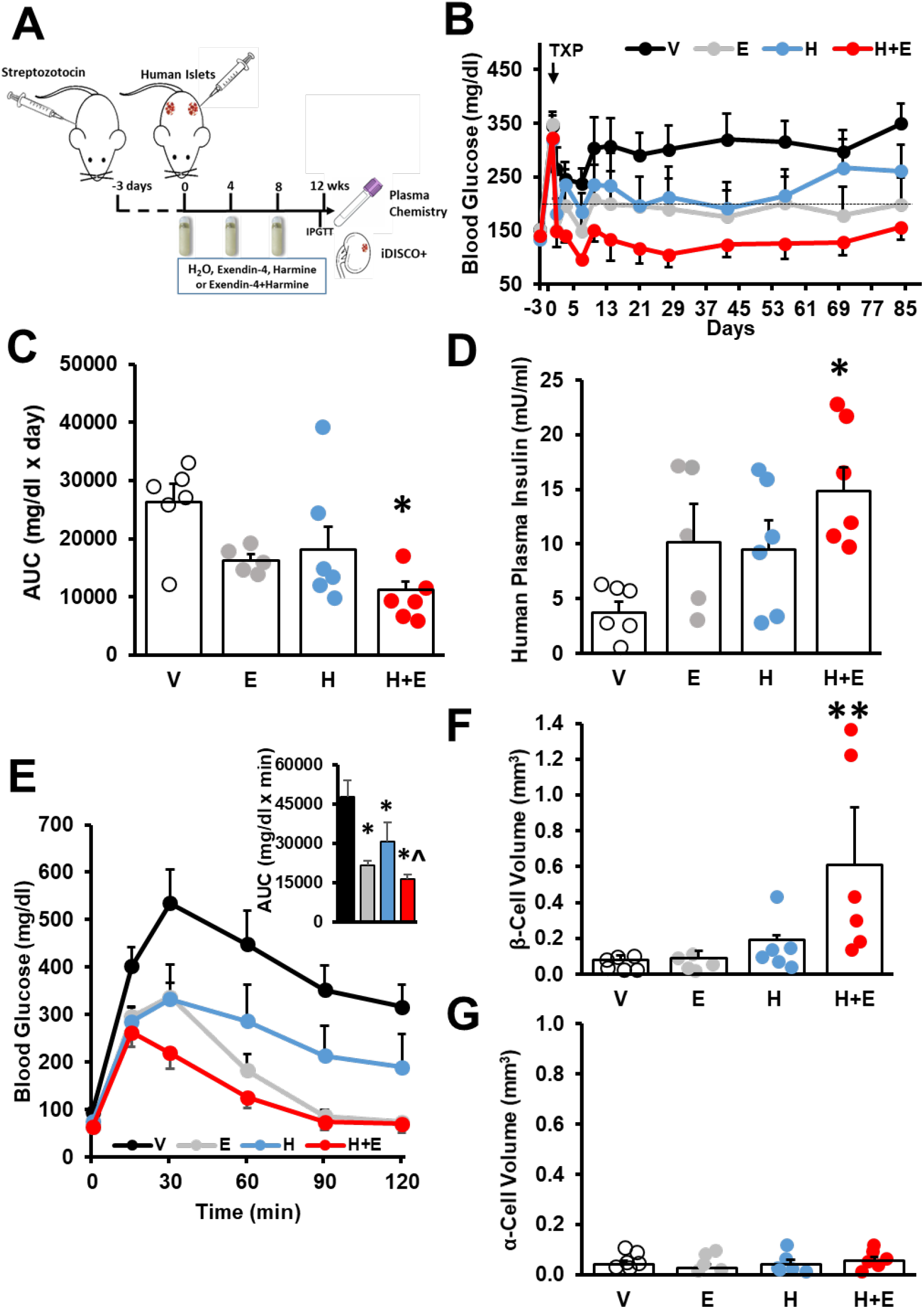
Effects Of Three Months Of Harmine and/or Exendin-4 On Human Beta Cell Mass, Function and Glucose Homeostasis In Streptozotocin-Induced Diabetic Mice. (**A**) The three-month study protocol in diabetic mice. Mice were rendered diabetic by streptozotocin injection and then 200-250 human IE per kidney were transplanted under the kidney capsule. Four-week minipumps were implanted and replaced at one month and two months to provide three months of continuous infusion. At month 3, kidneys were harvested and blood obtained. (**B**) Morning blood glucose at the time points indicated, before and after islet transplantation and during the three months of treatment, and (**C**) the area under the curve for the blood glucose levels for the four groups. Dots represent individual islet grafts from five to six different human islet donors in five to six different mice. (**D**) Plasma human insulin concentrations in the four groups at the end of three-month treatment. (**E**) Intraperitoneal glucose testing in the four groups. Inset shows area under the curve for each group. (**F**) Human beta cell volume and (**G**) human alpha cell volume in the four groups of mice at three months. In all panels, error bars indicate mean ± SEM, * indicates p<0.05 vs V, ** indicates p<0.05 vs V and E groups, ^ indicates p<0.05 vs V and H groups.

Notably, in this beta cell-deficient STZ model, human beta cell volume was significantly higher (~600%) in the harmine-exendin-4 group than the other three groups (**Fig. 10F**) and comparable to the 700% changes observed in the euglycemic group described earlier (**Fig. 4C**). Equally importantly, as had been observed in the euglycemic model (**Fig. 6I,J**), human alpha cell volume did not change in the STZ-diabetic model over three months, despite the marked increase in beta cell volume (**Fig. 10F,G**). Taken together, these data indicate that harmine-exendin-4 combination therapy expands human beta cell mass, and normalizes blood glucose levels in diabetic mice, and does so without altering alpha cell mass.

## Discussion

Here we show that DYRK1A inhibitors in combination with a canonical and widely employed GLP1RA lead to an increase in human beta cell mass, function and glycemic control *in vivo.* These findings apply both to human beta cells transplanted into euglycemic mice as well as into mice with severe diabetes. These observations should provide novel pharmacologic approaches that are scalable to millions of people with diabetes. This possibility raises a number of important questions. For example, how much beta cell regeneration and mass expansion is enough for T1D and T2D? Answering this can only be addressed through human trials evaluating C-peptide secretion, insulin reserve and glycemic control, but a 4-to 7-fold increase over three months in non-diabetic and, more importantly, in diabetic conditions is very encouraging. Specifically, to reverse the 50-60% reduction in beta cell mass in people with T2D (6,7), a 4-to 7-fold increase in three months would seem more than sufficient, and the accompanying improvement in beta cell differentiation and function should be particularly beneficial for T2D. T1D provides a greater challenge because baseline beta cell mass is lower in established T1D than in T2D (1,5), but a 4-to 7-fold increase in 3 months should improve glycemic control and reduce “fragility” and, of course, longer term treatment is feasible if it were needed. Beta cell autoimmunity will likely remain a challenge in T1D, but recent advances in the control of autoimmunity in T1D provide reason for optimism (38–40).

Since DYRK1A is ubiquitously expressed, one must worry about undesired off-target effects in tissues outside the beta cell. Safety and efficacy data at one-week (22) and at three-months (described herein) are encouraging, but will need to be assessed over even longer treatment periods, and in large animal preclinical studies before human safety and efficacy trials can be performed. Of note, many GLP1RA’s and DPP-IV inhibitors that elevate endogenous GLP1 are currently in use in millions of people with diabetes (36,41–42); simply adding an orally administered DYRK1A inhibitor in this setting to restore beta cell mass would be a simple, inexpensive and highly scalable approach to diabetes treatment. Optimizing dosing and defining clinically acceptable circulating concentrations of harmine and GLP1RA’s will need to be accomplished in subsequent human studies. Here, the breadth of GLP1RA’s and DPP-IV inhibitors available (36,41–43) and the flexibility in DYRK1A inhibitor dosing required for human beta cell proliferation (22) are promising: lower doses of DYRK1A inhibitors and GLP1RAs are certainly feasible. Some would argue that beta cell-specific targeting may be required for DYRK1A inhibitors, but again, the existing one-week (22) and the three-month safety data described herein suggest that such targeting may be unnecessary.

This current study takes advantage of our prior observation that low doses of harmine have a synergistic effect with GLP-1RAs to induce proliferation in cells expressing the GLP-1 receptor (22). For the current study, we selected a dose of harmine that increases beta cell proliferation *in vivo* but does not increase alpha cell proliferation, with the goal of expanding human beta cells without altering alpha cell mass. Gratifyingly, the co-treatment with low-dose of harmine and exendin-4 led to a remarkable increase in human beta cell proliferation and mass expansion in the absence of alpha cell proliferation at one-week or four-weeks of treatment, and only modest alpha cell proliferation at three months, resulting in no significant increase in alpha cell mass during the three month treatment. Importantly, no differences in circulating glucagon concentrations were observed. We attribute this beta cell selectivity to the low dose of harmine employed and the very low abundance or absence of GLP-1Rs on alpha cells (37).

The 4-to 7-fold increase in human beta cell mass over three months is surprising in the face of beta cell proliferation rates or labeling indices that are consistently higher than in controls, but are nonetheless low, below 1.0%. While this bodes well from a safety standpoint, it raises a question regarding the mechanism(s) responsible for the robust increase in human beta cell mass. Notably, in addition to beta cell proliferation, we find that beta cell survival in the immediate post-transplant period is enhanced, as evidenced by the lower rates of TUNEL labeling in the drug-treated groups; by greater beta cell mass at 1 week in the harmine-exendin-4 group than the other groups; and, by a smaller increase (3-fold) in the one month pre-transplant experiment (**Fig. 4J** vs. **Fig. 4G**) in which transplanted islets were allowed to engraft and vascularize before treatment initiation. This beta cell prosurvival effect appeared to be related to harmine and not to exendin-4, since both harmine alone and the combination consistently decreased human beta cell death in all the human islet preparations transplanted, while the effects of exendin-4 were variable and not significant. Interestingly, in this regard, harmine treatment *in vivo* has been shown to protect neurons and renal tubular cells from ischemic injury through potential antioxidant mechanisms (44–46).

The marked expansion of beta cell mass in the face of low beta cell proliferation rates, raises an additional intriguing possibility. Alpha cell proliferation rates at four weeks and three months were higher in the harmine and harmine-exendin-4 groups, yet no increase in alpha cell mass was observed (**Fig. 6**). We speculate that the harmine and harmine-exendin-4 combination may encourage alpha-to-beta cell transdifferentiation as has been reported in rodent islets (43,47–50). Documenting this possibility will require lineage tracing genetic studies which have been performed in rodent models using rodent alpha cell-specific gene promoters. Unfortunately, and despite great effort, no laboratory has developed reliable human alpha cell-specific lineage tracing tools. Until this occurs, a potential role for alpha-to-beta transdifferentiation must remain speculative.

One additional achievement is noteworthy. Accurate assessment of human beta cell mass in animal transplant models has been tedious and poorly reproducible or quantifiable. Thus, the novel marriage of iDISCO+ tissue clearing with 3D imaging - widely used in neuroscience (51–53) - to equally widely used models of human islet transplant allows accurate *in vivo* beta cell mass quantification to be widely performed in the diabetes research community for the first time. This provides a new technical standard in beta cell regeneration research. This tool should further accelerate the already rapid progress in beta cell regeneration and replacement research.

Despite remarkable new insights, this study has limitations. As noted above, the mechanisms underlying the marked increase in human beta cell mass remain incompletely defined, and would benefit from lineage tracing studies if and when genetic tools for such studies can be developed in the future for human alpha cells. Also, it remains uncertain whether beta cell-specific targeting will be necessary, and if so, what the ideal targeting molecule might be. Our preclinical histopathology and proliferation studies suggest that targeting may not be necessary, but longer term studies in higher preclinical models will be required to clarify this issue. Another important question relates to the durability of the 7-fold increase in human beta cell mass: will it decline when therapy is removed? Will it continue to increase with administration lasting more than three months? And will markers of proliferation decline to baseline when drug treatment is complete? Answering these questions will require extensive additional studies over the next several years.

In summary, we demonstrate that the combination of a DYRK1A inhibitor with a GLP1 receptor agonist markedly and rapidly enhances human beta cell proliferation, survival, beta cell mass expansion, and beta cell function over a very brief period of time. This appears to be acceptably beta cell-specific, avoiding off-target adverse effects on non-beta cell tissues and cell types, and maintains beta cell differentiation and function, both in a euglycemic model as well as in the conventional beta cell depletion model of diabetes. This approach would appear to be far less expensive and far more scalable to millions of people with diabetes, as compared to alternate approaches to restoring human beta cell mass, exemplified by pancreas or islet transplantation, or beta cell replacement through stem cell strategies. Further refinements in dosing, duration of treatment, reversibility as described above, and deeper exploration of safety and potential toxicities are required. Finally, we provide a novel and remarkably precise tool for assessing human beta cell mass in the form of 3-D imaging.

## Methods

### Human pancreatic islets

Human adult pancreatic islets from 26 non-diabetic donors were provided by the Integrated Islet Distribution Network and Prodo Laboratories. The average donor age was 46.5±2.4 and 57.7% were from male donors. Additional details are provided in **Supplemental Table 1**.

### Chemicals

Harmine-HCl was synthesized in the Drug Discovery Institute at Mount Sinai (54,55). Exendin-4 was purchased from MedChemExpress (Monmouth Junction, NJ).

### Human Islet Transplantation into RAG-1^−/−^ Immunodeficient Mice

Human islets were transplanted into the renal subcapsular space as described in detail previously (20–22). Numbers of human islet equivalents (IEQ, one IEQ = 125 μm diameter islet) are described in the Figure Legends. Islets were transplanted in 4-5 month-old normal euglycemic or streptozotocin-induced (200mg/Kg) diabetic RAG1^−/−^ mice as indicated in the text and figure legends.

### Minipump Delivery of Harmine and Exendin-4

Harmine and exendin-4 dissolved in water were loaded into Alzet (Cupertino, CA) model 1004 mini-osmotic pumps at a concentration of 27 mg/ml and 1 mg/ml, respectively, to permit subcutaneous delivery of harmine and exendin-4 for one month at a continuous rate of 3 mg/kg/day and 0.1 mg/kg/day, respectively. For three-month studies, pumps were replaced at 28 days and 56 days with new pumps and fresh harmine and exendin-4. For one week studies, the same type of pumps, dilutions and procedures were used. For the experiments in **Supplemental Fig. 1**, 2-3-month old male C57BL6/N mice (Charles River) were injected daily ip with 10mg/kg/day of harmine or minipumps were implanted to deliver vehicle, 1, 3, 10 mg/kg/day of harmine for one week.

### Human Islet Graft Collection

The kidneys containing the grafts were preserved by *in vivo* intracardiac perfusion in anesthetized mice with heparinized saline buffer (pH 7.4) followed by 4% paraformaldehyde (PFA; Electron Microscopy Sciences, Hatfield, PA, USA), and then immersed for 24 hours at 4 °C in the same fixative agent. For immunohistochemical analysis of beta cell proliferation (see below), right kidneys containing the islet grafts were transferred to 10% neutral buffered formalin before being embedded in paraffin. 5-μm-thick sections were cut and mounted on glass slides. For whole-mount staining and volume imaging, left kidneys containing the human islet grafts were washed three times with PBS before proceeding with the optical clearing protocols.

### Optical Clearing of Kidneys

Whole-mount staining and clearing was performed using a modified version of the iDISCO+ method (https://idisco.info/) termed iDISCO+ (51–53). In brief, kidneys were dehydrated at room temperature (RT) with gradual addition of methanol in PBS (20%, 40%, 60%, 80%, 100%), incubated in 100% dichloromethane (DCM) (Sigma-Aldrich, St. Louis, MO, USA) to remove hydrophobic lipids. Kidneys were bleached overnight with a 1:5 ratio of hydrogen peroxidase (H2O2):methanol at 4°C to reduce tissue autofluorescence. Tissues were rehydrated with a methanol gradient (80%, 60%, 40%, 20%) and incubated in permeabilization buffer with 5% DMSO/0.3 M Glycine in 0.1% Triton X-100/0.05% Tween-20/0.0002% heparin/0.02% NaN3 (PTxwH) in 1x PBS for 24 hours. Kidney were incubated in blocking buffer consisting of PTxwH with 3% normal donkey serum (Jackson Immunoresearch, West Grove, PA, USA) at 37°C for 48 hours. Primary antibodies [anti-insulin (A5064, Agilent, Santa Clara, CA), anti-glucagon (ab137817, Abcam, Cambridge, MA) and anti-smooth muscle actin (A5228, Millipore-Sigma, St. Louis, MO)] were diluted in blocking buffer and applied for 4-6 days at 37°C. Excess primary antibodies were washed for 24 hours with PtxwH with periodic solution changes that each lasted between 1-3 hours. Dye-conjugated secondary antibodies were diluted in blocking buffer (1:500) and samples incubated for 4 days at 37°C with rocking. Samples were washed 24 hours in PtxwH at 37°C with periodic solution changes. After five 60-min washes in PBS at RT, kidneys were dehydrated with a methanol gradient in PBS (20%, 40%, 60%, 80%, 100%). Samples were rinsed 3 times for 30 min in 100% methanol, followed by 3 washes 30-min each in dichloromethane (DCM, Sigma-Aldrich) for extraction of remaining lipids, before being transferred to dibenzyl ether (DBE, Sigma-Aldrich) for optical clearing. Samples were kept in DBE until the time of the tissue imaging analysis.

### Imaging Processing and Analysis of Kidneys

To obtain a 3D image of human beta cells transplanted into the kidney capsule, kidney samples were scanned continuously through an automatic 3D axis sample stage, and the fluorescence signals were detected by an objective that provides a series of 2D z-stack images. Subsequently, the acquired 2D z-stack images were analyzed by 3D image reconstruction. Z-stacked optical sections of the whole kidney were acquired with an Ultramicroscope II (LaVision BioTec, Bielefeld, Germany) at a 1.3x or 4x magnification. Laser intensities were set to prevent saturated pixels (typically 50-70% of maximum power). We used the following laser excitation wave lengths: 488 nm (Alexa 488) and 647 nm (Alexa 647). Z-stacked optical sections were acquired at 1.3x magnification with a 3 μm step size, or at 4x with a 3 μm step size and dynamic focus with a maximum projection filter. Whole kidneys were imaged at 1.3x with dynamic focus, and with multiple z-stacks acquired at 4x with a 20% overlap and tiled using the plugin TeraStitcher through the ImSpector Pro software (LaVision Biotec). Digital visualization of volume and surface images covering the insulin-positive islets to automatically determine volumes and intensity data were performed using Imaris Software version 9.1-9.3.1 (Bitplane, Belfast, United Kingdom). Volume reconstructions were performed using the surface function with absolute intensity as the threshold for detection of beta cells. A smoothing factor of 10 μm was used for islets analyzed at 1.3x, and a factor of 3.25 μm was used at 4x.

### Blood Glucose, Human Insulin, Glucagon, Exendin-4 and Harmine Measurements and Glucose Tolerance Test

An intraperitoneal glucose tolerance test (2 mg/kg) was performed on day 84 as previously described (20). Blood was obtained by tail vein nicking. Glucose was measured using AlphaTrak2 glucometer (Abbott, Alameda, CA). Plasma human insulin was measured using the human insulin ELISA kit (Mercodia, Winston Salem, NC) and plasma human glucagon was measured by the human glucagon ELISA kit (Thermo Scientific, Frederick, MD). Exendin-4 plasma levels were measured using the exendin-4 EIA kit (Phoenix pharmaceuticals, Burlingame, CA). Harmine was measured in plasma by liquid chromatography–mass spectrometry analysis by WuXi AppTec (Cranbury, NJ). Briefly, 20μL of plasma per mouse was protein precipitated with 200μL of a solution containing 100 ng/mL Labetalol and 100 ng/mL Tolbutamide as recovery standards in acetonitrile. The mixture was vortexed and then centrifuged at 4000 rpm for 15 min at 4°C. An aliquot of 100 μL of the supernatant was transferred to sample plate and mixed with 100 μL water, then the plate was shaken at 800 rpm for 10 min and 2 μL supernatant was then injected for LC-MS/MS in a Triple Quad™ 6500+ LC-MS/MS System (Sciex, Framingham, MA).

### Human Islet Graft Immunohistochemistry

At the end of the treatment, animals were sacrificed, right kidneys containing the islet graft were harvested, fixed paraffin-embedded and analysis of alpha and beta cell proliferation was performed in sections immunolabeled with anti-insulin (A5064, Agilent, Santa Clara, CA or 6N-ID4, Developmental Studies Hybridoma Bank, DSHB, Iowa City, IA) or anti-glucagon (ab137817, Abcam, Cambridge, MA) and anti-Ki67 (MA5-14520, Thermo Scientific, Waltham, MA) antibodies and DAPI, as previously described (20–22). Labeled cells were then visualized using fluorescent phase contrast microscopy (Zeiss Axiovert Vert, White Plains, NY). Human alpha and beta cell proliferation was quantified as Ki67+ and insulin+ or glucagon+ cells and 1,000-2,000 alpha or beta cells were blindly counted per sample. Beta cell death was determined in islet graft sections stained for insulin or glucagon using the terminal deoxynucleotidyl transferase-mediated dUTP nick end-labeling (TUNEL) method (Promega, Madison, WI) (20) and at least 1,000 beta cells were blindly counted per sample. Islet graft sections were also immunolabeled with antibodies for PDX-1 (ab47308, Abcam), MAF-A (ab239356, Abcam) and NKX6.1 (F55A10-C, DSHB) and insulin (as above) as previously described (20–22). Beta cell size was analyzed in insulin stained islet grafts sections as previously described (56), and the size of at least 300 beta cells was quantified per section using ImageJ (National Institutes of Health, Bethesda, MD).

### Tissue Histopathology

Tissues were collected after intracardiac perfusion as indicated above and then immersed for 24 hours at 4 °C in 4% paraformaldehyde. Liver, intestine, pancreas, spleen, adipose tissue, kidney, heart and lung were transferred to 10% neutral buffered formalin before being embedded in paraffin. 5-μm-thick sections were cut and mounted on glass slides. Sections were stained for hematoxylin and eosin for double-blinded pathology analysis. Consecutive mouse tissue sections were immunolabeled for Ki67 (550909, BD Pharmingen) and actin (A2066, Sigma), Ki67 (550909, BD Pharmingen) and cytokeratin-19 (CK-19) (ab52625, Abcam) and Ki67 (MA5-14520, Thermo Scientific), insulin (GTx39371, Genetex, Irvine, CA) and glucagon (G2654, Sigma) as detailed previously (20–22). At least 1,000 cells per field were counted. Brains were sectioned on a vibrating microtome (50μm). Free-floating sections were washed three times in PBS for 15 minutes followed by 1-hour incubation at RT in blocking solution. Sections were then incubated for 72 hours in primary antibody diluted in blocking solution at 4°C. The sections were then washed in PBS three times followed by a 2-hour incubation in secondary antibody diluted in blocking solution at RT. After a further series of three wash steps, sections were mounted (Fluoromount with DAPI, Southern Biotech) and covered. The final concentrations of the primary and secondary antibodies and blocking solutions used in the brain sections are as follows: Rabbit anti-cFos (Cell signaling 2250S), donkey anti-rabbit 647 (Jackson ImmunoResearch 711-605-152) were diluted in 3% donkey serum, 0.1% Triton X in PBS. Rabbit anti-Ki67 (Thermofisher RM-9106-S1), donkey anti-rabbit 647 (Jackson ImmunoResearch 711-605-152) were diluted in 10% donkey serum, 0.3% Triton-X in PBS. Images were acquired as Z-stacks using a Zeiss Inverted LSM 780 laser scanning confocal microscope (Carl Zeiss MicroImaging, Inc.). For quantification purposes, exposure times were set based on the control group and were comparable among the different experimental groups. Immunolabeled cells were quantified using FIJI software. The investigator and the pathologist were both blinded to treatment group. Every third section was collected, stained, and imaged.

### Statistics

Data presented as bar graphs and scatterplots show means ± SE. Statistical analysis was performed using one-way ANOVA with Tukey or Holm post hoc HSD (http://astatsa.com) for comparison among groups. P ≤ 0.05 was considered statistically significant.

### Study Approval

All protocols were performed with the approval of and in accordance with guidelines established by the Icahn School of Medicine at Mount Sinai Institutional Animal Care and Use Committee.

## Author contributions

A.G.-O., S.A.S., R.J.D, and A.F.S. conceived of the studies, oversaw the experiments and wrote the manuscript. C.R., A.A., P.W., Y.L., R.L., K.B., H.L., G.L., V.G., N.T., and K.K. performed the studies.

## Acknowledgements

We thank the Bonnie and Joel Bergstein family and the Lonnie and Thomas Schwartz families for their constant support. We thank the JDRF, the NIDDK-supported Einstein-Sinai DRC and the Human Adenovirus and Islet Core, The Mount Sinai Imaging Core for light sheet microscopy use and support, the NIDDK Integrated Islet Distribution Program and Prodo Labs for supplying human cadaveric islets. This work was supported by JDRF grant 2-SRA-2017 514-S-B, NIH grants P-30 DK020541, R-01 DK116873, R-01 DK116904, R-01 DK125285, R-01 DK105015, R-01 DK113079, R-01 DK126450, R-01 NS097184 and OT2 OD024912, NSF grant 1930157, DoD grant W81XWH-20-1-0345, the American Diabetes Association Pathway to Stop Diabetes Grant 1-17-ACE-31, a Mindich Child Health and Development Institute Pilot and Feasibility Grant and an Icahn School of Medicine Distinguished Scholar Award. A.A. is supported by a postdoctoral fellowship from the Charles H. Revson Foundation and Swedish Society for Medical Research (SSMF).

## Data Availability

All data are available on request from the Corresponding Author.

