## Supplemental Information for "Human Beta Cell Mass Expansion In Vivo With A Harmine and Exendin-4 Combination: Quantification and Visualization By iDISCO+ 3D Imaging"

**The Harmine and Exendin-4 Combination Selectively and Markedly Enhances Human Beta Cell Mass in Vivo.**

**Supplemental Figure 1.**

**Figure Legend of Supplemental Figure 1.**

**Figure Legends of Supplemental Videos 1 to 3.**

**Supplemental Videos 1 to 3.**

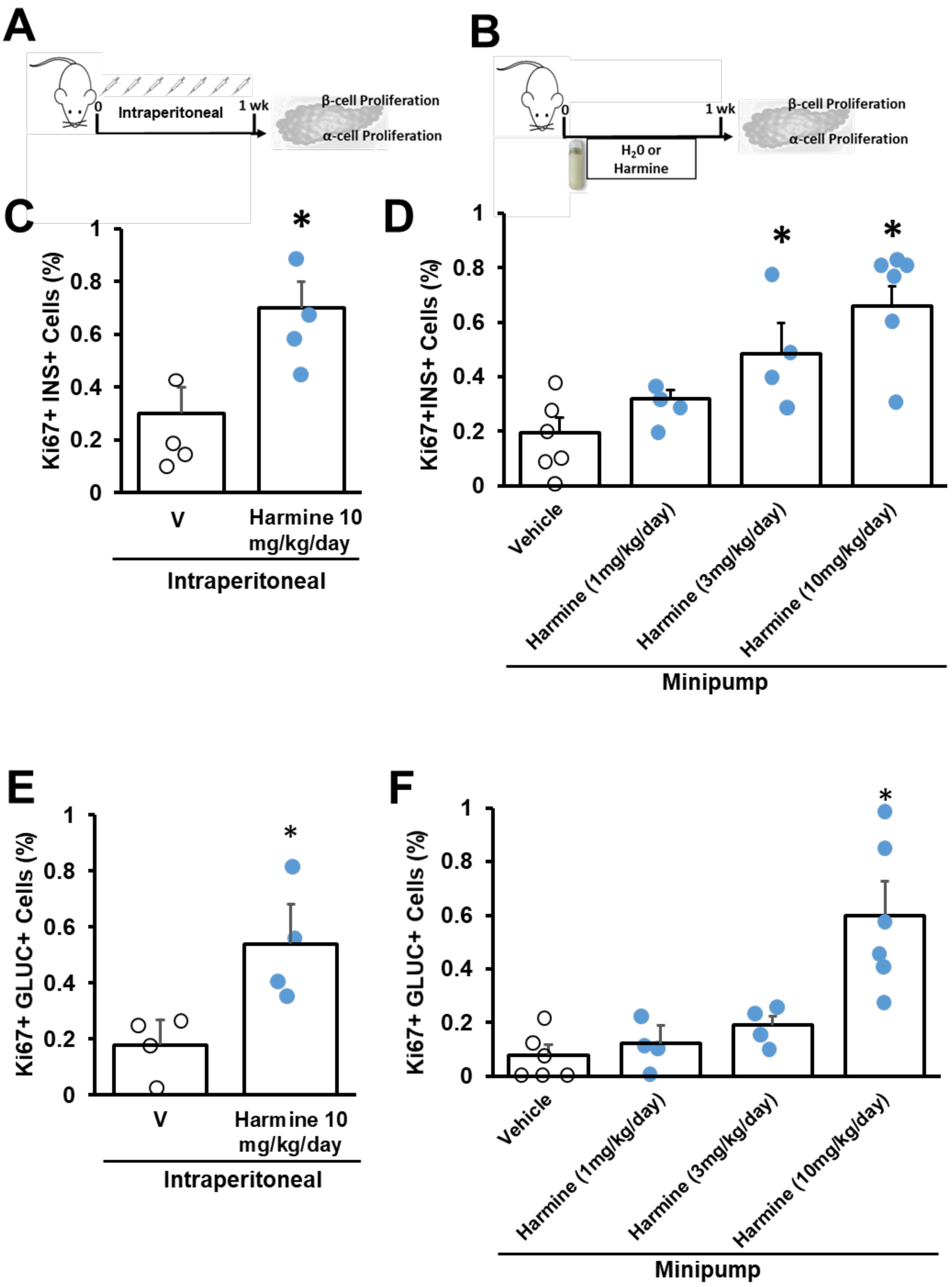


**Supplemental Figure 1. Effects Of Daily Intraperitoneal Or Continuous Infusion Of Harmine For One Week On Mouse Alpha and Beta Cell Proliferation In C57BL6N mice.** (**A**) The one week study protocol in mice. Mice were either injected daily intraperitoneally with vehicle or 10mg/Kg of harmine. In another set of mice, minipumps were implanted for delivery of vehicle or 1, 3 and 10mg/Kg/day of harmine. At one week, pancreata were harvested and alpha and beta cell proliferation was measured. Labeling index of Ki67 in (**B**) beta cells and (**C**) alpha cells in 4-6 different mice per treatment. Dots represent each individual mouse. In all panels, error bars indicate mean ± SEM, * indicates p<0.05 vs V.

**Supplemental Video 1.** iDISCO+ immunolabeling of a whole mouse kidney containing a human islet graft. (A) Movie shows the kidney immunolabeled for insulin (green) and smooth muscle actin (red).

**Supplemental Video 2.** iDISCO+ immunolabeling of a whole mouse kidney containing a human islet graft from a mouse treated with vehicle for three months. Movie shows the kidney immunolabeled for insulin (green) with volume rendering of the images using Imaris. This video relates to Fig. 4B.

**Supplemental Video 3.** iDISCO+ immunolabeling of a whole mouse kidney containing a human islet graft from a mouse treated with harmine plus exendin-4 (H+E) for three months. Movie shows the kidney immunolabeled for insulin (green) with volume rendering of the images using Imaris. This video relates to Fig. 4B.
