## Supplemental Table 1 for "Human Beta Cell Mass Expansion In Vivo With A Harmine and Exendin-4 Combination: Quantification and Visualization By iDISCO+ 3D Imaging"

| Human islet preparation | 1 | 2 | 3 | 4 | 5 | 6 | 7 | 8 | 9 | 10 | 11 | 12 | 13 |
|---|---|---|---|---|---|---|---|---|---|---|---|---|---|
| MANDATORY INFORMATION | | | | | | | | | | | | | |
| Unique identifier | HP-18277 | HP-18330 | HP-18336 | HP-19003 | HP-19017 | HP-19030 | HP-19047 | HP-19073 | HP-19163 | HP-19189 | HP-19192 | HP-19199 | HP-19219 |
| Donor age (years) | 62 | 53 | 61 | 63 | 45 | 25 | 53 | 50 | 24 | 37 | 62 | 59 | 41 |
| Donor sex (M/F) | M | F | F | F | F | M | F | M | M | M | F | F | M |
| Donor BMI ($kg/m^2$) | 35.9 | 26.3 | 24.5 | 27 | 36.1 | 27.3 | 22.7 | 21.8 | 24.5 | 32.2 | 30.1 | 24.7 | 35.1 |
| Donor $HbA_{1c}$ or other measure of blood glucose control | 5.20% | 5.40% | 5.30% | 5.80% | 5.70% | 5.20% | 4.80% | 5.50% | 5.60% | 5.20% | 5.20% | 5.00% | 5.80% |
| Origin/source of islets[b] | PRODO | PRODO | PRODO | PRODO | PRODO | PRODO | PRODO | PRODO | PRODO | PRODO | PRODO | PRODO | PRODO |
| Islet isolation center | PRODO, Aliso Viejo, CA | PRODO, Aliso Viejo, CA | PRODO, Aliso Viejo, CA | PRODO, Aliso Viejo, CA | PRODO, Aliso Viejo, CA | PRODO, Aliso Viejo, CA | PRODO, Aliso Viejo, CA | PRODO, Aliso Viejo, CA | PRODO, Aliso Viejo, CA | PRODO, Aliso Viejo, CA | PRODO, Aliso Viejo, CA | PRODO, Aliso Viejo, CA | PRODO, Aliso Viejo, CA |
| Donor history of diabetes? Yes/No | NO | NO | NO | NO | NO | NO | NO | NO | NO | NO | NO | NO | NO |
| RECOMMENDED INFORMATION | | | | | | | | | | | | | |
| Donor cause of death | Stroke | Stroke | Stroke | Stroke | Stroke | Anoxic event | Stroke | Head Trauma | Anoxic event | Head Trauma | Stroke | Stroke | Head Trauma |
| Warm ischaemia time (h) | Unkown | Unkown | Unkown | Unkown | Unkown | Unkown | Unkown | Unkown | Unkown | Unkown | Unkown | Unkown | Unkown |
| Cold ischaemia time (h) | Unkown | Unkown | Unkown | Unkown | Unkown | Unkown | Unkown | Unkown | Unkown | Unkown | Unkown | Unkown | Unkown |
| Estimated purity (%) | 90% | 80% | 90% | 85% | 90% | 90% | 90% | 90% | 95% | 85% | 80% | 90-95% | 90% |
| Estimated viability (%) | 95% | 95% | 95% | 95% | 95% | 95% | 95% | 95% | 95% | 95% | 90% | 95% | 95% |
| Total culture time (days)[d] | 8 | 4 | 5 | 7 | 7 | 7 | 7 | 6 | 7 | 4 | 6 | 8 | 7 |
| Glucose-stimulated insulin secretion or other functional measurement[e] | Unkown | Unkown | Unkown | Unkown | Unkown | Unkown | Unkown | Unkown | Unkown | Unkown | Unkown | Unkown | Unkown |
| Handpicked to purity? Yes/No | Yes | Yes | Yes | Yes | Yes | Yes | Yes | Yes | Yes | Yes | Yes | Yes | Yes |
| Additional notes | | | | | | | | | | | | | |

| | | | | | | | | | | | | |
|---|---|---|---|---|---|---|---|---|---|---|---|---|
| 14 | 15 | 16 | 17 | 18 | 19 | 20 | 21 | 22 | 23 | 24 | 25 | 26 |
| HP-19226 | HP-19242 | HP-19265 | HP-19276 | HP-19324 | HP-19333 | Samn13149867 (HU1181) | HP-20010 | Samn13938639 (HU1188) | HP-20191 | HU1197 | SAMN15656747 | SAMN15770453 |
| 51 | 55 | 61 | 37 | 43 | 38 | 28 | 58 | 50 | 39 | 44 | 24 | 48 |
| M | M | M | M | F | M | M | M | F | F | M | M | F |
| 31.3 | 28.4 | 33.1 | 23.9 | 22.8 | 24.9 | 30.8 | 23.2 | 39.2 | 22.1 | 28.84 | 33.6 | 30.9 |
| 5.20% | 5.60% | 4.50% | 5.80% | 5.40% | 5.30% | 5.20% | 5.70% | 5.00% | 5.60% | 5.50% | Unkown | Unkown |
| PRODO | PRODO | PRODO | PRODO | PRODO | PRODO | IIDP | PRODO | IIDP | PRODO | IIDP | IIDP | IIDP |
| PRODO, Aliso Viejo, CA | PRODO, Aliso Viejo, CA | PRODO, Aliso Viejo, CA | PRODO, Aliso Viejo, CA | PRODO, Aliso Viejo, CA | PRODO, Aliso Viejo, CA | Southern California Islet Cell Resources Center | PRODO, Aliso Viejo, CA | Southern California Islet Cell Resources Center | PRODO, Aliso Viejo, CA | Southern California Islet Cell Resources Center | University of Wisconsin | University of Wisconsin |
| NO | NO | NO | NO | NO | NO | NO | NO | NO | NO | NO | NO | NO |
| Anoxic event | Stroke | Stroke | Head Trauma | Stroke | Head Trauma | Head Trauma | Stroke | Stroke | Head Trauma | Stroke | Head Trauma | Stroke |
| Unkown | Unkown | Unkown | Unkown | Unkown | Unkown | 11mins | Unkown | Unkown | Unkown | Unkown | 0 | Unkown |
| Unkown | Unkown | Unkown | Unkown | Unkowm | Unkown | 8.4 | Unkown | 11.6 | Unkowm | 7.13 | 6.53 | 328 mins |
| 90% | 90-95% | 90% | 90% | 95% | 85% | Unkown | 90% | 80% | 95% | 80% | 92% | 90% |
| 95% | 95% | 95% | 95% | 95% | 95% | Unkown | 95% | 95% | 95% | Unkowm | 99% | 99% |
| 5 | 6 | 5 | 6 | 7 | 4 | 4 | 4 | 3 | 6 | 3 | 4 | 5 |
| Unkown | Unkown | Unkown | Unkown | Unkown | Unkown | Unkown | Unkown | Unkown | Unkown | Unkown | Unkown | Unkown |
| Yes | Yes | Yes | Yes | Yes | Yes | Yes | Yes | Yes | Yes | Yes | Yes | Yes |
